# A cell culture platform for *cryptosporidium* that enables long-term cultivation and new tools for the systematic investigation of its biology

**DOI:** 10.1101/134270

**Authors:** Christopher N. Miller, Lyne Jossé, Ian Brown, Ben Blakeman, Jane Povey, Lyto Yiangou, Mark Price, Jindrich Cinatl, Wei-Feng Xue, Martin Michaelis, Anastasios D. Tsaousis

**Affiliations:** Laboratory of Molecular & Evolutionary Parasitology, RAPID group, School of Biosciences, University of Kent, Canterbury, UK; School of Biosciences, University of Kent, Canterbury, UK; Industrial Biotechnology Centre, School of Biosciences, University of Kent, Canterbury, UK; School of Physical Sciences, University of Kent, Canterbury, UK; Institut für Medizinische Virologie, Klinikum der Goethe-Universität, Frankfurt am Main, Germany

## Abstract

*Cryptosporidium* parasites are a major cause of diarrhoea that pose a particular threat to children in developing areas and immunocompromised individuals. Curative therapies and vaccines are lacking. Currently, *Cryptosporidium* oocysts for research must be freshly produced in animals and cannot be long-term stored. Here, we show that COLO-680N cells infected with two different *Cryptosporidium parvum* strains (Moredun, Iowa) produce sufficient infectious oocysts to infect subsequent cultures. Oocyst identity was confirmed by specific staining (Crypt-a-glo, Vicia Villosa lectin, Sporo-glo), PCR-based amplification of *Cryptosporidium*-specific genes, lipidomics fingerprinting, and atomic force microscopy (AFM). Antibody-stained oocysts produced unstained oocysts confirming production of novel oocysts. Infected cultures could be cryoconserved and continued to produce infectious oocysts after resuscitation. Transmission electron microscopy identified all key *Cryptosporidium* life cycle stages. Infected cultures produced thick-walled (primarily involved in *Cryptosporidium* transmission between organisms) and thin-walled oocysts (important for *Cryptosporidium* propagation within a host/tissue) as indicated by DAPI staining (only thin-walled oocysts are permeable to DAPI staining, thus allowing visualisation of sporozoites) and AFM. In conclusion, we present a novel, easy-to-handle cell culture system that enables the propagation, cryopreservation and detailed investigation of *Cryptosporidium* at a laboratory scale. Its availability will accelerate research on *Cryptosporidium* and the development of anti-*Cryptosporidium* drugs.

---

Cryptosporidiosis causes a significant number of deaths in children and immunocompromised individuals^1-4^. It is caused by species of the genus *Cryptosporidium*, in humans typically by *C. parvum* and *C. hominis*. The *Cryptosporidium* species belong to the phylum Apicomplexa and it has recently been proposed for the species to be reclassified as a member of the subclass of gregarina^5, 6^. They are waterborne pathogens, and cryptosporidiosis has commonly been associated with disease in developing countries^7-9^. However, more recent molecular epidemiology studies suggested that the disease is also an increasing health concern in developed countries and may have reached epidemic levels^1, 2, 10-14^. Only one moderately effective drug (nitazoxanide) is available for the treatment of cryptosporidiosis. More effective drugs are urgently needed^10, 15, 16^.

*Cryptosporidium* is a parasite that invades host cells, within the boundaries of the host cell membrane, residing intracellularly yet extra-cytoplasmic sometimes referred to simply as epicellular^17^. *Cryptosporidium* typically infects epithelial tissues of the upper intestinal tract, accompanied by localised deterioration of microvilli (**Supplementary Fig. S1** displays the life cycle of the parasite20). In immunocompromised individuals, the parasite can also be found in other epithelial tissues, including most of the upper stages of the digestive and respiratory tracts as well as other unrelated organ systems organs^18, 19^. The *Cryptosporidium* life cycle is complex and involves a number of intracellular/extracytoplasmic and extracellular stages resulting in oocysts that contain the infectious sporozoites^20^.

A cell culture system that enables the continuous *Cryptosporidium* cultivation and systematic elucidation of the *Cryptosporidium* life cycle^11^, especially the endogenous phases^21^, is missing. Previous approaches were hampered by problems including rapid senescence of primary cell cultures, incomplete parasite life cycles, and insufficient production of sporulated infectious oocysts^10, 11, 18, 22, 23^. The current methods used to produce infectious *Cryptosporidium* oocysts, aside from small-scale cultures *in vitro*, require continuous infection of animals, typically neonatal cows or sheep and sometimes mice^11, 24^. Due to a lack of cryopreservation methods, oocysts cannot be stored and need to be freshly prepared on a continuous basis. A recent publication tackled the challenge of cell culture-based oocyst production using a hollow fiber technology that mimics the gut^25^. However, specialised equipment is needed and the required cell culture media supplements are expensive. In addition, the system does not enable the studying of the *Cryptosporidium* life cycle and biology in real time at a cellular level in the context of a host cell.

Here, we show that inoculation of COLO-680N cultures with *C. parvum* produced sufficient amounts of infectious oocysts to enable the sustainable propagation of the parasite in standard tissue culture at a laboratory scale.

## Results

### COLO-680N cell cultures enable the continuous and sustainable propagation of *Cryptosporidium parvum.*

We tested a panel of seven human cancer cell lines for their capacity to support *C. parvum* propagation including COLO-680N (oesophageal squamous-cell carcinoma), DLD-1 (colon adenocarcinoma), KYSE-30 (oesophageal squamous-cell carcinoma), HCT-15 (colorectal adenocarcinoma), SJSA-1 (osteosarcoma), MKN-1 (gastric carcinoma), and the colon adenoma carcinoma cell line HCT-8, which has most commonly been used for the investigation of *Cryptosporidium* in cell culture^10, 26^. However, *Cryptosporidium*-infected HCT-8 cultures do not produce enough infectious oocysts to maintain infected cultures^11, 27^, which also raise concerns about the suitability of HCT-8 for the studying of *Cryptosporidium* biology.

The cell lines were infected with the *C. parvum* strain Moredun^28, 29^ using an input of 50,000 of excysted oocysts per mL. After an incubation period of two weeks, COLO-680N cultures were the only ones that had produced substantially more oocysts (approx. 75-fold) higher than the number of input oocysts (**Fig. 1a** and **Supplementary Table S1**). While HCT-8 cells had died after 14 days of infection, COLO-680N cultures remained viable and produced oocysts for almost eight weeks without sub-culturing, requiring only weekly media exchange (**Fig. 1b**). As a result, total *Cryptosporidium* oocyst production in the COLO-680N cell line (number of produced oocysts) exceeded the HCT-8-mediated oocyst production (2.5 x 10^5^ oocysts/ mL) by 20 times (5 x 10^6^) after ten days of incubation (**Fig. 1c**). At day 60, COLO-680N cells had produced an accumulated number of 1.2 x 10^7^ oocysts/ mL obtained from weekly harvests. Notably, oocysts derived from the supernatants of COLO-680N cell cultures, but not from the supernatants of HCT-8 cell cultures, enabled the infection of novel cell cultures (**Supplementary Fig. S6c**). Infection of COLO-680N cells with cattle-derived *C. parvum* addition, we also performed three rounds of infection using COLO-680N culture-derived oocysts without noticing changes in oocyst production efficacy showing that COLO-680N cells are suited for the continuous long-term cultivation of *C. parvum* oocysts.

**Figure 1:**
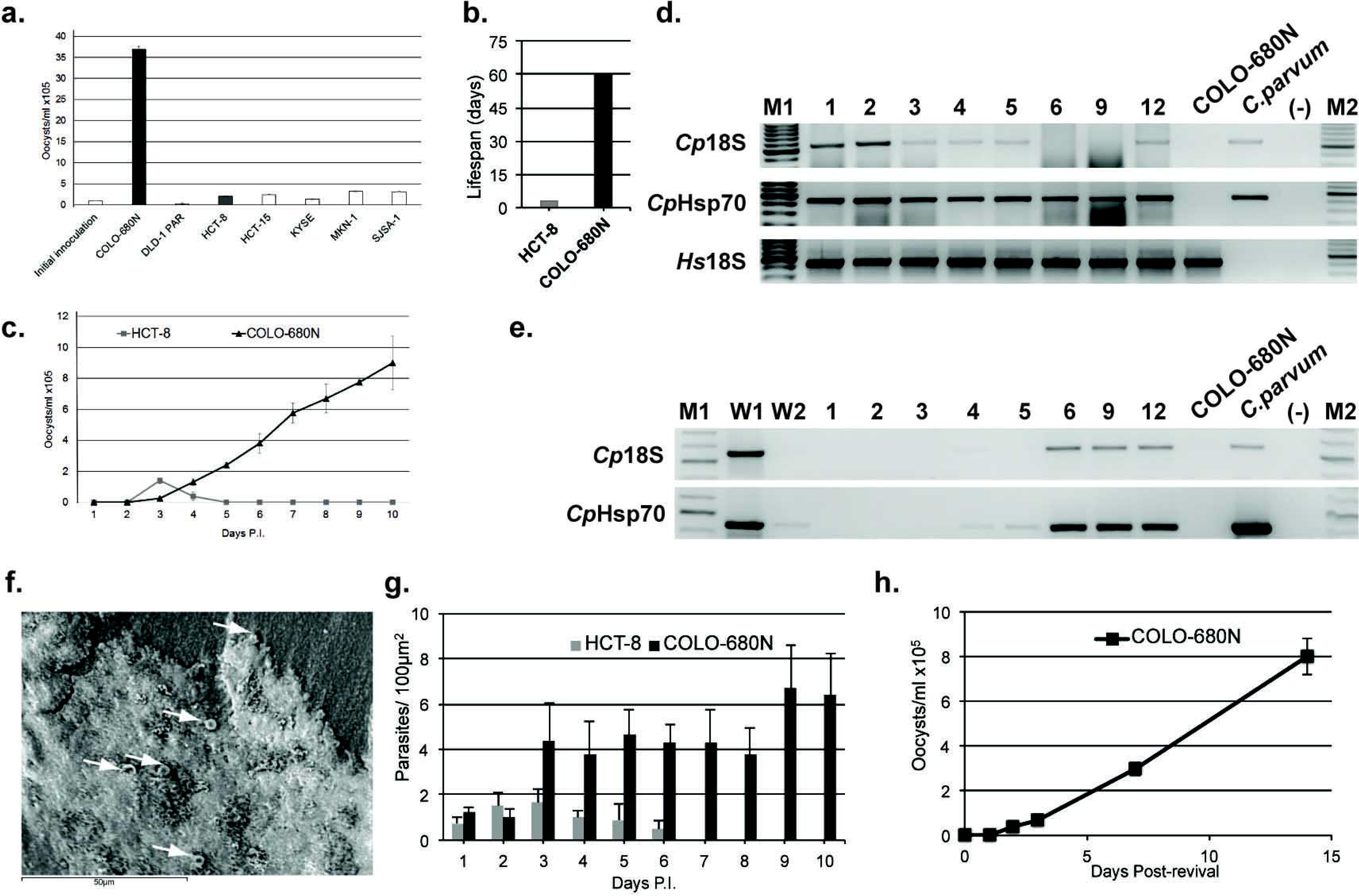
Cell culture-based production of *C. parvum* oocysts. A bar chart representing the average *C. parvum* oocyst production (mean ± S.D. from three independent experiments) in the investigated cell lines after initial infection with 1 x 10^5^ excysted oocysts. Final oocyst counts are representative of total content recovered after 14 days of incubation, regardless of host cell viability. Initial experiments returned a near 30-fold return in oocysts by COLO-680N cultures, compared to only a 2-fold return by HCT-8. **b.** Bar chart of the time span during which oocysts were produced by COLO-680N and HCT-8 after a single initial inoculation (mean ± S.D. from three independent experiments). **c.** *C. parvum* oocyst production in COLO-680N and HCT-8 cancer cells over a 10-day period after an inoculation with 1 x 10^5^ excysted oocysts, measured through daily sampling (mean ± S.D. from three independent experiments). **d.** PCR amplification of *C. parvum* 18sRNA (*Cp*18S, 580 bp, primers CF/CR) and Hsp70 (*Cp*Hsp70, 462 bp, primers Hsp70F4/Hsp70R4) DNA fragments from *C. parvum*-infected COLO-608N cells. A *Homo sapiens* 18s DNA fragment (*Hs*18s, 418 bp, primers Hs18s1F/Hs18s1R) demonstrates abundance of host cell DNA in the sample. DNA extraction was performed at day 1, 2, 3, 4, 5, 6, 9 and 12-post infection. Cattle-derived *C. parvum* oocysts (*C. parvum*) and uninfected COLO-680N cells (COLO-680N) served as controls. M1 is the 1 Kbp DNA ladder from Promega. M2 is the 100 bp DNA ladder from Promega. **e.** PCR amplification of *C. parvum* 18S RNA (*Cp*18S, 580 bp, primers CF/CR) and Hsp70 (*Cp*Hsp70, 462 bp, primers Hsp70F4/Hsp70R4) DNA fragments from samples derived from the supernatants of *C. parvum*-infected COLO-608N cells via percoll gradient after an excystation protocol. Input oocysts were removed by two washing steps with PBS (W1 & W2) 6 hours post infection, leaving no detectable *C. parvum* DNA in suspension. Time points and controls were the same as described in (d.). **f.** Scanning electron microscopy of COLO-680N produced *C. parvum* oocysts. White arrows indicate *Cryptosporidium* oocysts. **g.** Bar chart demonstrating the average number of *C. parvum*-infected cells in a 100 μm^2^ oil field at 1000x magnification at day 1 to 10 post infection (mean ± S.D. from five independent experiments). Parasites were identified as the presence of co-localised propidium iodide and Sporo-glo within a host cell. **h.** Oocyst production in *C. parvum*-infected COLO-680N cell cultures after two weeks of cryopreservation and resuscitation (mean ± S.D. from three independent experiments).

*C. parvum* infection of COLO-680N cells was further confirmed using PCR to detect intracellular *C. parvum* DNA, indicating the continuous presence of *C. parvum* (**Fig. 1d**), and *C. parvum* DNA in the supernatants, indicating *C. parvum* extracellular life stages and oocysts production (**Fig. 1e**). *Cryptosporidium-*specific primers did not produce bands in non-infected COLO-680N cells (**Fig. 1d,e** and **Supplementary Fig. S2**). The amplified DNA regions were sequenced to confirm their identity. In addition, purified COLO-680N-produced oocysts were visualised by scanning electron microscopy (**Fig. 1f**).

The identity of the COLO-680N-produced *C. parvum* oocysts was further confirmed using different specific staining methods. Crypt-a-glo (Waterborne™; an antibody that recognises the oocyst cell wall), Vicia Villosa lectin (VVL, Vector labs; binds to specific O-glycan mucin repeats on *C. parvum* sporozoites), a mucin-like glycoprotein that contains a C-type lectin domain (CpClec; binds to surface of the apical region and to dense granules of sporozoites and merozoites^30^) and direct sporozoite staining using propidium iodide and Sporo-glo (Waterborne™) resulted in virtually identical staining patterns in *C. parvum*-infected COLO-680N cells indicating the presence of oocysts and other non-extracellular life stages of *Cryptosporidium* (**Fig. 2a and Supplementary Fig. S3, S4, S6 & S11**). Crypt-a-glo staining did not reveal any differences between COLO-680N-and cattle-produced oocysts (**Fig. 2e**). It has been reported that *Cryptosporidium* infections results in the production of thick and thin oocysts^31^. Thick-walled oocysts are thought to be primarily involved in *Cryptosporidium* transmission from one organism to another whereas thin oocysts may play an important role in the maintenance and propagation of a *Cryptosporidium* infection in a given host or tissue^20^. Double staining of COLO-680N-produced *C. parvum* oocysts by Crypt-a-glo (stains the wall of thick and thin oocysts) and DAPI (can only penetrate thin-walled oocysts, but not thick-walled oocysts) suggest the presence of both types of oocysts (**Fig. 2d; Supplementary Fig. S11**). The comparison of Crypt-a-glo staining of *C. parvum*-infected COLO680N-and HCT-8 cells further confirmed that *C. parvum-*infected COLO-680N cultures are characterised by enhanced numbers of infected cells compared to *C. parvum*-infected HCT-8 cultures (**Fig. 1g** and **Supplementary Fig. S3)**. To finally confirm the production of fresh oocysts, Crypt-a-glo stained oocysts were excysted (**Supplementary Fig. S5a**) and used for the infection of COLO-680N cultures. Then, cell cultures were washed to remove remaining Crypt-a-glo stained oocysts. Upon harvesting neither the infected cultures nor the newly produced oocysts displayed Crypt-a-glo staining. However, oocysts were detected using DAPI indicating that indeed new oocysts were produced (**Supplementary Fig. S5**). We have also subsequently been able to propagate successfully the altenative *C. parvum* Iowa strain in COLO-680N cells (**Supplementary Fig. S6**).

**Figure 2:**
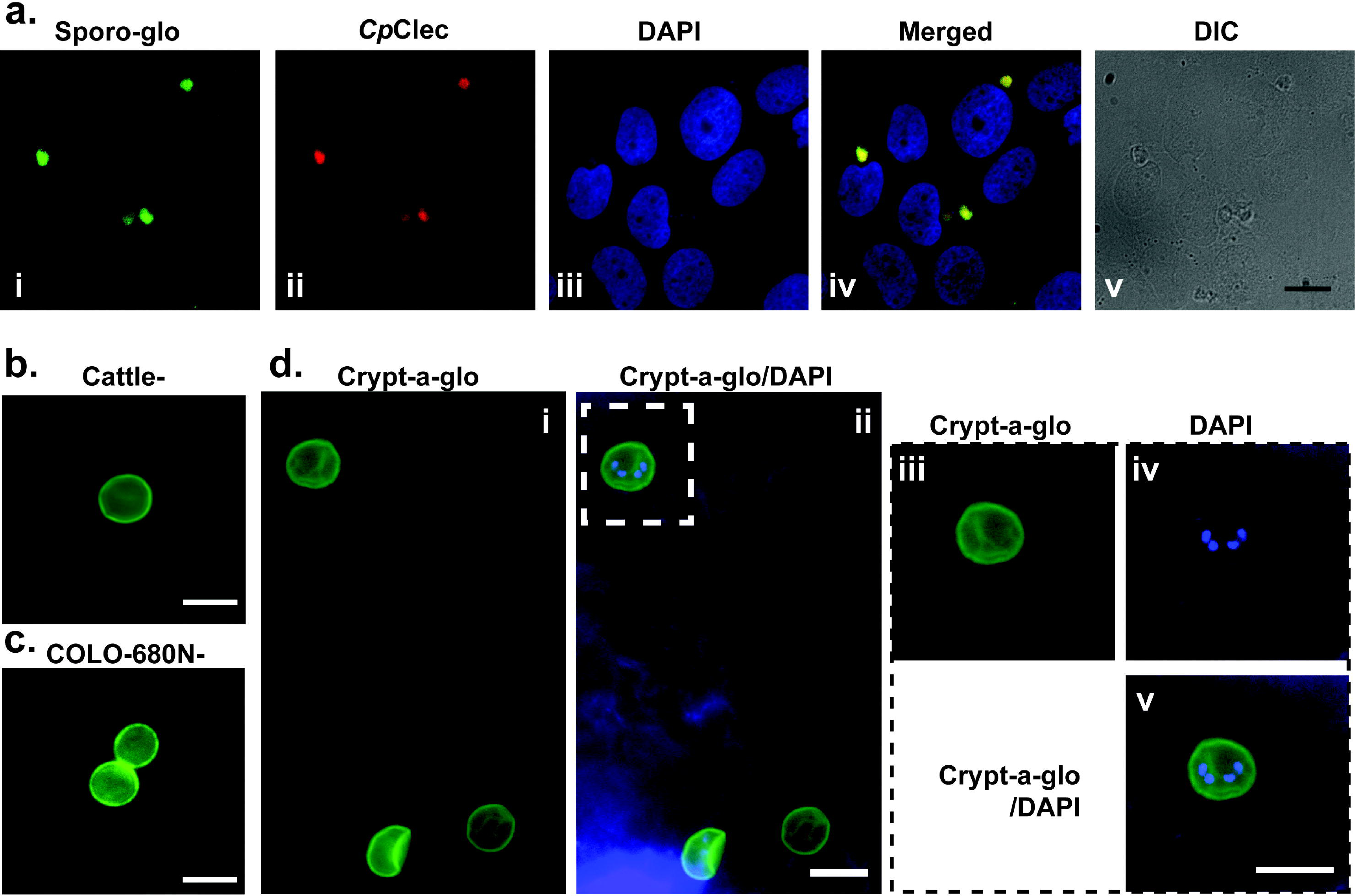
Detection of *C. parvum* using different specific staining methods. **a.** Visualisation of *C. parvum* oocysts in infected COLO-680N cells. *C. parvum* oocysts were detected using Sporo-glo (Waterborne), a fluorescein-labelled mouse monoclonal antibody binding to *Cryptosporidium* (**i**), CpClec, that binds to the surface of the apical region and to dense granules of sporozoites and merozoites^30^ (**ii**), and DAPI staining (**iii**) that can be used to distinguish between host cell nuclei and parasites by morphological inference when coupled with DIC and other stains. **iv.** Merge of **i, ii and iii** conclusively showing that what is being observed is indeed *C. parvum* oocysts. **v.** The corresponding differential interference contrast microscopy image. Scale bar 40 μm. **b.** Crypt-a-glo-stained cattle-produced oocyst. Scale bar 5 μm. **c.** Crypt-a-glo-stained COLO-680N-produced oocysts. Scale bar 5 μm. **d.** COLO-680N-produced oocysts stained with Crypt-a-glo and DAPI to differentiate between thick and thin oocysts. DAPI can only penetrate thin oocysts resulting in staining of the sporozoites but not thick oocysts. Results indicate three oocysts, two thick oocysts that only show Crypt-a-glo staining and one thin oocyst that shows both Crypt-a-glow and DAPI staining (**i.** Crypt-a-glo, **ii.** DAPI, **iii.** merge). **iii-v.** Inset from **ii** showing the thin oocyst at higher magnification indicating DAPI staining of the four sporozoites. Scale bar 5 μm.

### Cryopreservation of *C. parvum*-infected COLO-680N cultures

Currently, there is a lack of a cryopreservation system that enables the long-term storage of infectious *Cryptosporidium* parasites^32^. Here, *C. parvum* strain Moredun-infected COLO-680N cells were cryopreserved, stored for two weeks at −80°C, and resuscitated by standard protocols used for cell cultures. Three days after resuscitation, the cultures started to produce oocysts in a similar fashion like freshly infected COLO-680N cultures (**Fig. 1h**). This demonstrates that *C. parvum*-infected COLO-680N can be cryoconserved, providing the first long-term storage system for *Cryptosporidium*.

### Mass spectrometry-based fingerprinting indicate differences between the proteomes of *C. parvum*-infected and non-infected COLO-680N cells, but not between infected and non-infected HCT-8 cells

Next, we compared *C. parvum*- and non-infected cell cultures by a MALDI-TOF mass spectrometry-based fingerprinting approach. Principal Component Analysis (PCA) analysis of the pre-processed data, as described in supplementary methods and in more detail in Povey *et al,* 2014^36^, resulted in separate groupings of the COLO-680N, but not the HCT-8 samples. (**Supplementary Fig. S7a)**. We found substantial alterations in the fingerprints between non-infected and *C. parvum*-infected COLO-680N cells, five days after infection, but not between non-infected and *C. parvum*-infected (**Supplementary Fig. S7b-e**). These findings suggest *C. parvum* infection resulted in a more noteworthy difference in COLO-680N cultures, compared to HCT-8, suggesting either a more successful infection (presence of an increase number of *Cryptosporidium*-originated proteins) or a more pronounced effect on the host cell proteome during infection.

### COLO-680N-and cattle-produced *C. parvum* oocysts are indistinguishable by lipidomics and atomic force microscopy (AFM)

Furthermore, we compared COLO-680N and cattle-produced *C. parvum* oocysts by a lipidomics approach and by atomic force microscopy (AFM). The lipidomics characterisation was performed using MALDI-TOF mass spectrometry for the analysis of lipids within the range of 600 to 2,000 daltons (**Supplementary Fig. S8a-d**). Graphical representation of the Principal Components (PC1 and PC2) from PCA^36^ showed groupings consistent with samples having a large degree of similarity between one another (**Supplementary Fig. S8e-f**).

To investigate the existence of oocysts in the highest magnification possible, we employed AFM that has been previously used to elucidate unique surface details at a level of resolution not visible using any other imaging modalities in other parasites (e.g. *Giardia* and *Trypanosoma*^37^). Notably, we observed two types of oocysts by AFM in *C. parvum*-infected COLO-680N cultures (**Fig. 3**). We found a larger type of COLO-680N-produced oocysts (**Supplementary Fig. S9a** and **Supplementary Fig. S10**) that was indistinguishable from cattle-produced oocysts by force-distance curve-based imaging (**Supplementary Fig. S9b**). These oocysts are likely to represent thick-walled oocysts, since the cattle-produced oocysts are expected to be thick-walled oocysts that transmit infection from organism to organism^31^. However, we also identified a smaller type that most likely represents thin-walled oocysts and that may contribute to the continuous *C. parvum* infection pattern that we observed in cell culture (**Fig. 3b**), since thin-walled oocysts are thought to be responsible for infection dissemination within organisms and tissues^20^.

**Figure 3:**
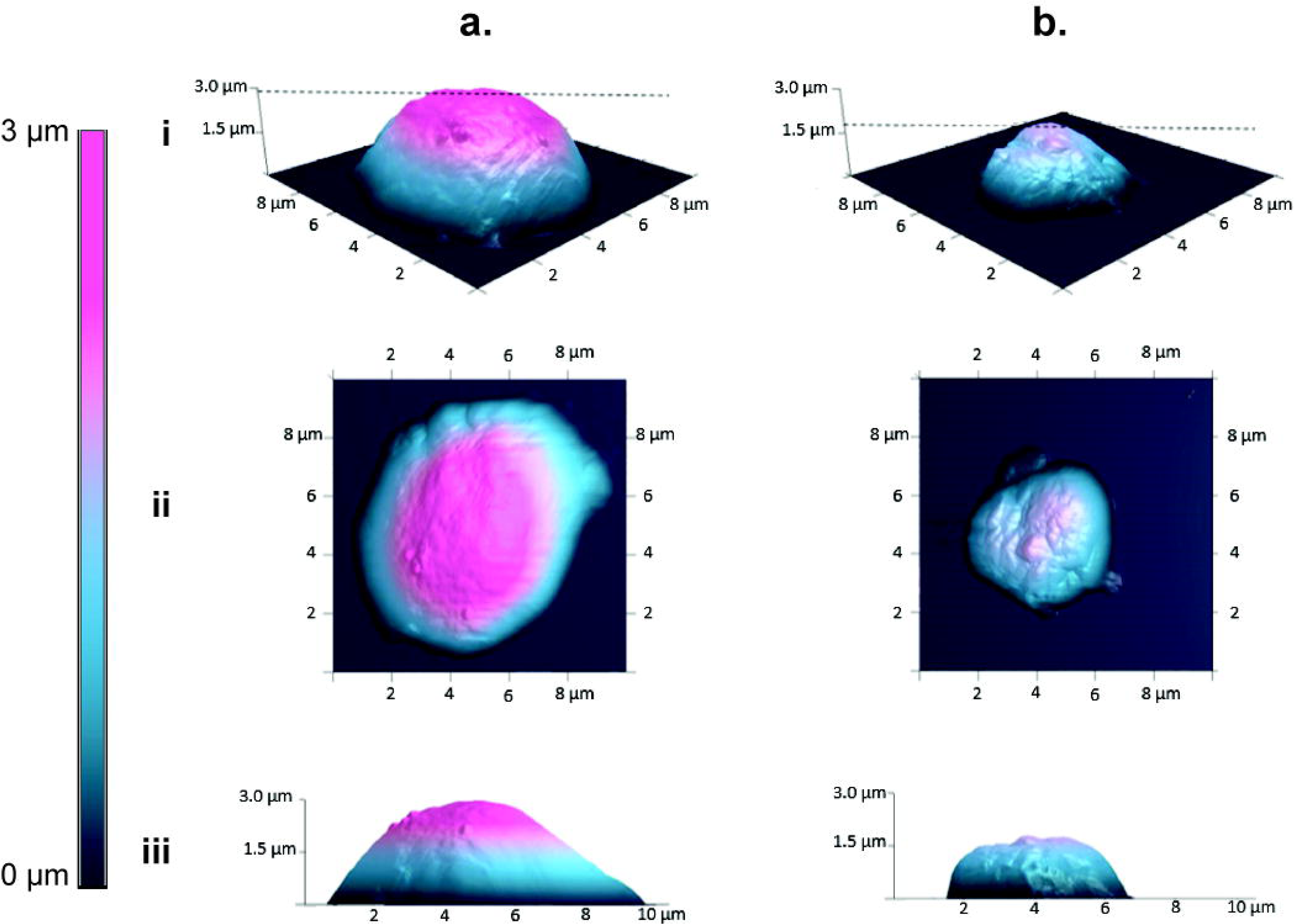
Oocysts captured by Atomic Force Microscopy (AFM) 3-D AFM height images displaying the overall morphology of thick (**a.**) and thin (**b.**) oocysts. (i) perspective view showing X, Y and Z dimensions (maximum height given by dotted line). Dimensions: Thick – (X) 9.5 μm, (Y) 7.5 μm and (Z) 3 μm; Thin – (X) 5 μm, (Y) 5 μm and (Z) 1 μm. (ii) top down view and (iii) side view of oocysts. All images are scaled to the same height contrast 0 – 3 μm (scale bar on left), which provides a visual aid in determining dimensional differences.

### Monitoring of *C. parvum* life cycle stages in COLO-680N cultures by transmission electron microscopy

Major stages of the *Cryptosporidium* life cycle including sporozoite, trophozoite, meront, macrogamont, microgamont, zygote, immature oocyst, and matured oocyst have been defined based on the pathological investigation of infected tissue^3, 18, 26, 28^. However, many *in vitro* studies report on a failure to detect all life cycle stages or the detection of abnormalities at particular phases of the *C. parvum* life cycle^38, 39^. After synchronising *C. parvum* infection in COLO-680N cells using a previously established protocol^40^, we could confirm the presence of the known key *C. parvum* life cycle stages by transmission electron microscopy (**Fig. 4**).

**Figure 4:**
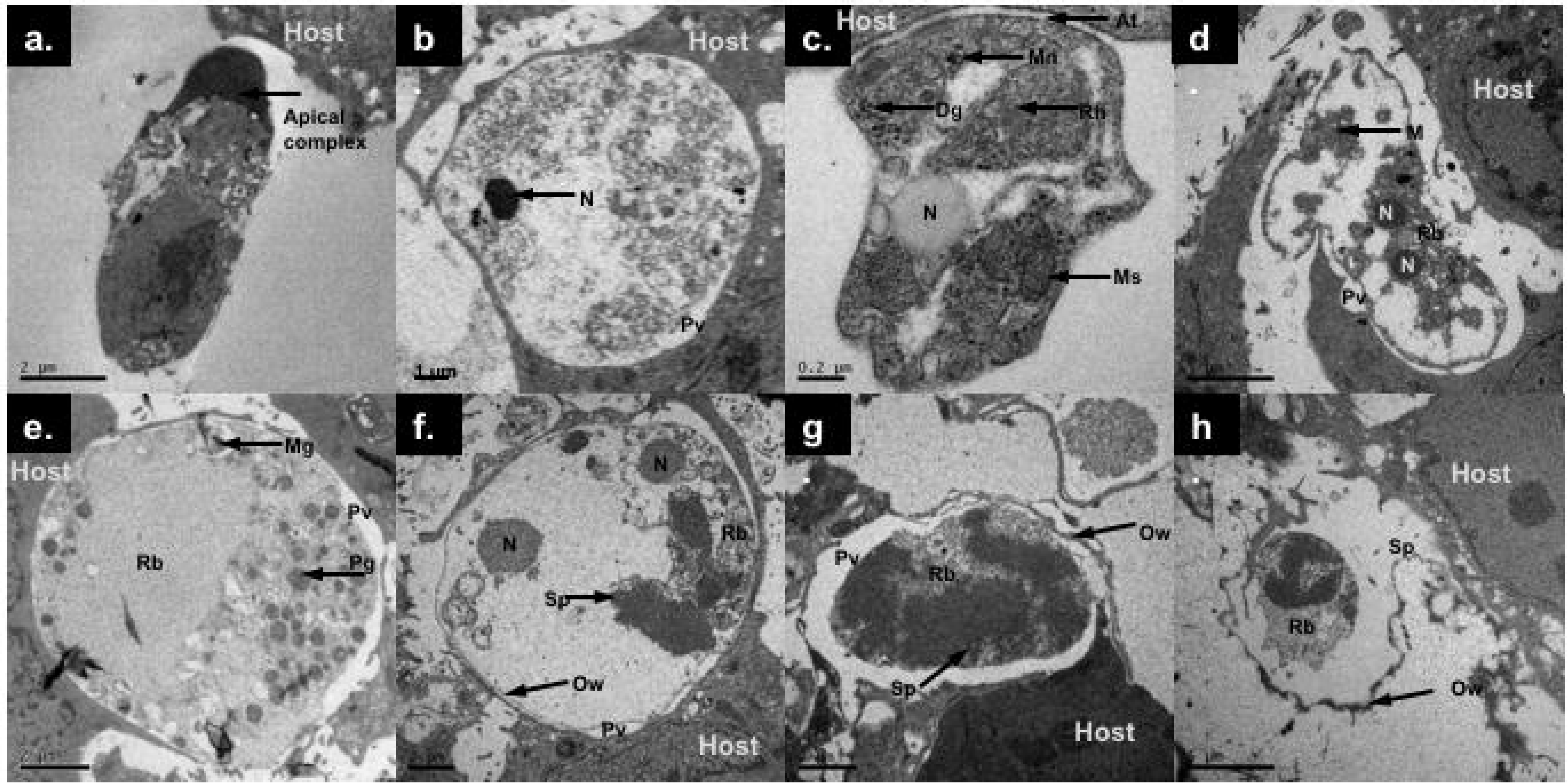
Visualisation of *Cryptosporidium* life-cycle progression using Transmission Electron Microscopy (TEM) Visualisation of *Cryptosporidium* life-cycle progression using Transmission Electron Microscopy (TEM). Images interpreted by visualising TEM images from published peer-reviewed literature^56-58^. TEM images of the main stages of *C. parvum* in COLO680N cells: **a.** sporozoite (sp) with visible apical complex; **b.** Early transition of a trophozoite/meront visible within a host cell, the nucleus (N) and parasitophorous vacuole (Pv) are discernible; **c.** a merozoite stage associates with the membrane of a COLO-680N cell (At), several organelles are visible including micronemes (Mn), Dense granules (Dg), a rod shaped rhoptry (Rh), Nucleus (N) and the mitosome (ms); **d.** a large meront with visible multiple nuclei (N), Remnant body (Rb) and a developing merozoite (M); **e.** a *C. parvum* gamont with characteristic polysaccharide granules (Pg) forming around a remnant body with a budding microgamete (Mg); **f.** a fertilised gamont with divided nuclei (N) and visibly thicker wall (Ow) shows early sporulation (Sp) from the remnant body (Rb); **g.** the majority of the zygote space is occupied by the remnant body (Rb) and sporozoites (Sp); **h.** an almost fully sporulated oocyst adhered to the cell surface, showing increased stress on the host cell membrane.

Moreover, we observed rare unique structures, apparently as a result of the interaction of the parasite with the host cell. For example, we could visualise the presence of the feeder organelle (**Fig. 5a**) that connects the host cell to the parasitophorous vacuole, as previously reported^41, 42^. Feeder organelles have been postulated to be involved in facilitating nutrient transport to the parasite, but had not been detected before in cell culture^43, 44^. Another interesting observation was the identification of the *Cryptosporidium* remnant mitochondrion (mitosome) and the crystalloid body in other life-stages than the sporozoites, including trophozoites, merozoites and microgamonts (**Fig. 5b-c**). Previously, both organelles had only been detected in *Cryptosporidium* oocysts or the excysted sporozoites^43, 45-47^. In addition, we observed the presence of dense arrangements of the host cytoskeleton in the periphery of the vacuole along with the development of the parasitophorous vacuole of *C. parvum* (**Fig. 5d**), suggesting a parasite-induced intracellular/epicellular rearrangement of host cells as supported by previous work^48, 49^.

**Figure 5:**
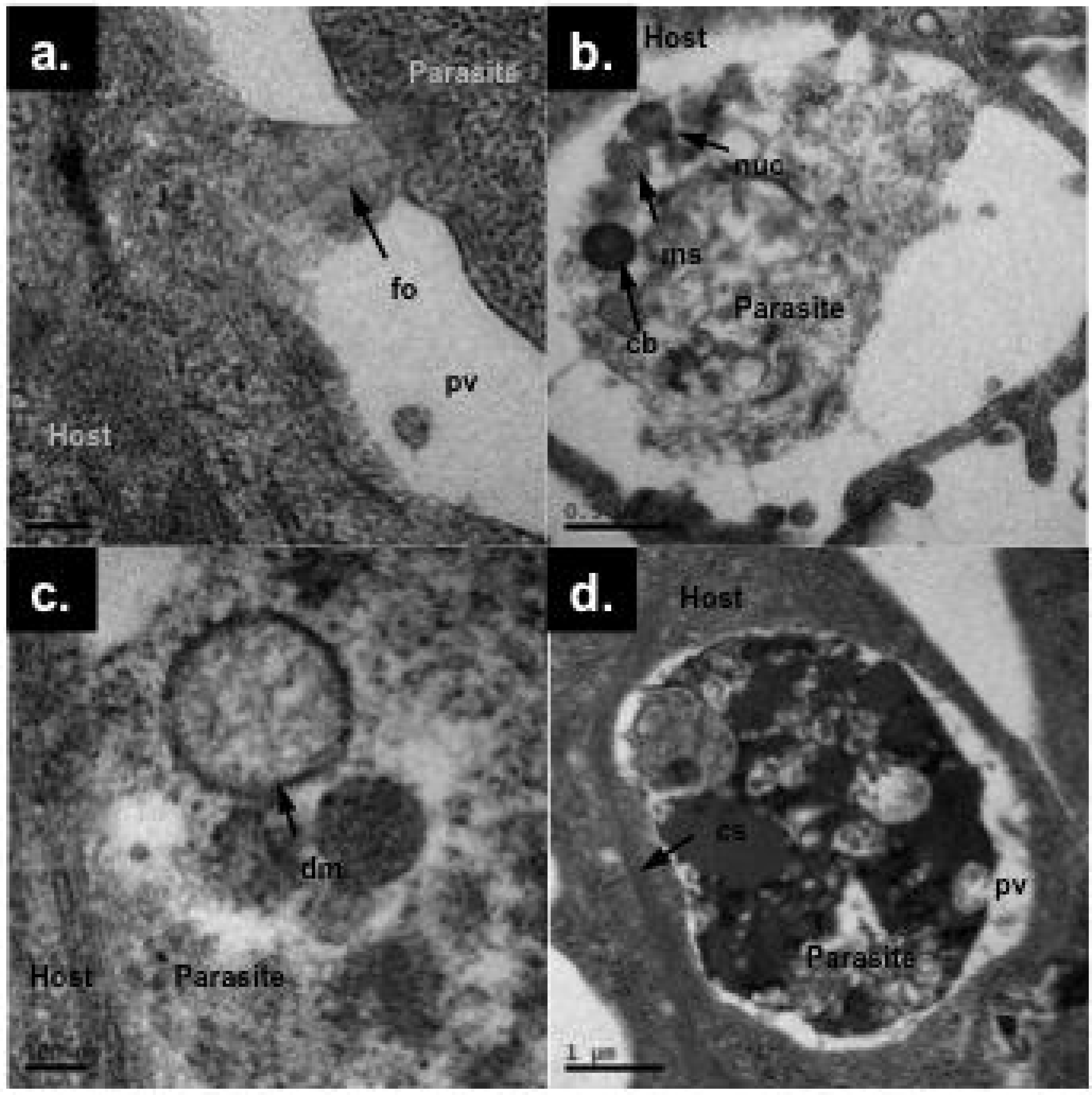
Unusual intracellular structures during *C. parvum* infection. TEM images showing the formation of numerous organelles and structures that are formed within the parasite and between the parasite and its host cell and not seen in other apicomplexan parasites. **a.** TEM image showing the formation of the feeder organelle (fo), which anchors the main body of the parasite to the peripheral of the parasitophorous vacuole (pv). **b.** TEM image showing numerous organelles within the sporozoite that has recently invaded a host cell. The electron dense structure that can be seen is the crystalloid body (cb). The nucleus (nuc) and the mitosome (ms) can also be seen in this image. **c.** TEM image demonstrating a clearly identifiable double membrane structure of the mitosome within a merozoite form of *C. parvum*. **d.** TEM image of *Cryptosporidium* epicellular stages showing the presence of dense arrangements of host cytoskeleton (cs) around the parasitophorous vacuole.

## Discussion

Here, we present a cell culture system that enables the sustainable, continuous propagation of infectious *C. parvum* oocysts and the systematic investigation of the *Cryptosporidium* life cycle. Previously attempts to cultivate *Cryptosporidium* in cell culture were affected by a lack of the production of sufficient amounts of infectious *C. parvum* oocysts^10, 11, 18, 22, 38, 39^ or required sophisticated, expensive specialist equipment and methodologies to support 3D cultures that are not commonly available to research laboratories ^25^. Moreover, 3D cultures do not enable the studying of the *C. parvum* biology^25^. In contrast, COLO-680N cells enable *C. parvum* propagation, the sustainable production of infectious *C. parvum* oocysts, and the investigation of the *C. parvum* biology at a laboratory scale in standard tissue cultures with commonly available equipment and know-how. In addition, these data demonstrate a long-term maintenance of the cell-line and subsequently a prolonged production of oocysts. The reasons behind this observation are unknown, but previous studies on COLO-680N have suggested that the expression of high-levels of fatty acid synthase might promote cell viability^33^. This could be beneficial to the parasite that depends on host cell lipid synthesis, since it is unable to synthesize fatty acids *de novo*^34, 35^.

We detected all key *C. parvum* life cycle stages in infected COLO-680N cells. Moreover, the presence of additional, previously unrecognised structures indicated that the systematic analysis of *C. parvum* replication in COLO-680N cells will provide novel insights resulting in a substantially improved understanding of the *C. parvum* biology. COLO-680N-produced oocysts were indistinguishable from cattle-produced oocysts by staining with antibodies that specifically bind to the *C. parvum* oocyst cell wall and by utilising lipidomics, and atomic force microscopy techniques. Thus, we are presenting for the first time, a new collection of tools (lipidomics and AFM) for investigating the cell biology and the composition of *Cryptosporidium,* which can be incorporated in further studies to provide a better understanding of the infection and life-cycle of the parasite.

It remains unclear why COLO-680N cells, in contrast to other cell lines that have been investigated so far, support *C. parvum* propagation. Nevertheless, our whole cell MALDI-ToF fingerprinting studies suggested that *C. parvum* infection of COLO-680N cells results in a substantial change in the features of the cultures, while *C. parvum* infection of HCT-8 cells, a model commonly used for the studying of *C. parvum,* did not^10, 39^. This may indicate a specific susceptibility of COLO-680N cells towards *C. parvum* infection. There also exists anecdotal evidence that suggests a link between squamous cell carcinomas and cryptosporidiosis which, given the nature of these results, we believe warrants closer examination^50^. In addition, we detected the presence of different types of oocysts (suggestive of the presence of thin oocysts) in COLO-680N cultures by Crypt-a-glo/ DAPI double staining and atomic force microscopy. The presence of thin-walled oocysts may also contribute to the successful *C. parvum* propagation, since thin-walled oocysts are thought to be responsible for the maintenance and dissemination of infection within tissues and organisms^20, 31^.

In conclusion, the discovery of COLO-680N as a cell culture platform for the production of *C. parvum* will provide a step-change with regard to research on *Cryptosporidium*: 1) It is the first easy-to-handle system that enables the long-term sustainable production of infectious oocysts at a laboratory scale and removes the constant dependence on immunosuppressed animals for production of *Cryptosporidium* oocysts along with all its ethical implications. 2) *C. parvum*-infected cell cultures can be frozen and stored. Prior to the establishment of the COLO-680N cultivation system for *C. parvum*, oocysts had to be freshly acquired from animals and could not be stored over longer periods. 3) Our study paves the way for establishment of compound-screening platforms for the identification of anti-*Cryptosporidium* drugs and the systematic elucidation of the *Cryptosporidium* biology, including the utilisation of a CRISPR transfection system for *Cryptosporidium*^24^. Hence, the COLO-680N-based platform for *C. parvum* propagation will enable a much greater community to work on *Cryptosporidium* and open unprecedented opportunities to decipher the *Cryptosporidium* biology and to develop anti-*Cryptosporidium* therapies.

## Methods

### Cell culture

The cell lines used in this study were: COLO-680N (Human oesophageal squamous-cell carcinoma), obtained from CLS Cell Line Services, Eppelheim, Germany; DLD-1 (Human colon adenocarcinoma), KYSE-30 (Human oesophageal squamous-cell carcinoma) and HCT-15 (human colorectal adenocarcinoma), obtained from DSMZ, Braunschweig, Germany; SJSA-1 (osteosarcoma) and HCT-8 (ileocecal colorectal adenocarcinoma), obtained from ATCC, Manassas, VA, US, Cat No. CRL-2098 and CCL-244, respectively; MKN-1 (gastric carcinoma), obtained from JCRB Cell Bank, Osaka, Japan (**Table S1**).

Cells were cultivated in RPMI-1640 medium (Sigma-Aldrich, Cat No R8758) supplemented with 10 % foetal bovine serum (Sigma-Aldrich, Cat No F8084), 100 U/mL penicillin, 100 μg/mL streptomycin and 250 ng/mL amphotericin B (Antibiotic Antimycotic solution, Sigma-Aldrich, Cat No A5955) at 37 °C and 5 % CO_2_.

### Excystation of C. parvum oocysts and infection of cell lines

*Cryptosporidium parvum* oocysts were provided by the Creative Science Company pre-purified and identified as the ‘Moredun’ strain, which was originally acquired from the gastrointestinal contents of a dead red deer calf (*Cervus elaphus*), sourced from the Glen Saugh experimental deer farm (Scotland, UK) in 1987^28, 29^. *Cryptosporidium parvum* oocysts Iowa strain were obtained from Bunch Grass Farm (Idaho, USA). For a typical infection, 1 x 10^5^ oocysts, counted prior to excystation, were used to infect T25 cell culture flasks at between 70-80 % confluency (2 x 10^5^ cells) giving a multiplicity of infection (MOI) of approximately 0.5. All infections were conducted in triplicate for reliability.

Excystation was achieved by adding 100 μL of 0.01% Trypsin and 400 μL of 0.5 % Sodium Hypochlorite to a pellet containing the desired number of oocysts^51, 52^. The homogenised mixture was then incubated in a 37°C water bath for one hour, with intermittent vortexing. The excystation procedure was monitored by phase contrast microscopy, using a haemocytometer and stopped when visible sporozoites exceeded 80 % of the theoretical maximum (4x the original number of oocysts present per sample). Samples were then pelleted at 2,200 x g for eight minutes, and suspended in cell culture medium, prior to cell infection. 24-26 h post-infection, T25 flasks were washed twice with 10 ml 1x PBS, to remove un-excysted oocysts and remaining sporozoites. Fresh media was then added. In order to assess the effectiveness of the post-infection washes, each wash was subjected to PCR testing (see below) for the presence of *C. parvum* DNA. A negative result would confirm that the only remaining *C. parvum* at this point would be intracellular stages, i.e. successful infections. Therefore any future positive PCR results would indicate the presence of a successful infection and production of oocysts after the initial infection (see **Fig. 1**). Enumeration of oocysts produced by the cultures was achieved via phase contrast microscopy of the purified oocysts fractions, using a standard haemocytometer. Where appropriate, estimates of infection intensity were also made using a fluorescence approach, where the number of *C. parvum* stages in a 1,000 x magnified field of view, identified by Sporo-glo or Crypt-a-glo, were averaged from 10 randomly selected areas of a stained culture. This method was used primarily in assessing the success of an infection for quality control purposes.

### Purification of Cryptosporidium from infected cultures

Growth media from infected cultures and a subsequent 5 mL wash (with 1x PBS) were collected. The suspensions were centrifuged at 500 x g for 5 minutes to remove host cells and debris. The supernatant, which contain the oocysts, was then transferred to fresh tubes, and the oocysts were pelleted by centrifugation at 2,100 x g for 8 minutes. The pellets were re-suspended in 1 mL of 1x PBS and carefully laid on top of 9 mL of saturated (37%) sodium chloride solution, in a 15 mL Falcon tube. These two layers were then topped up with 1 mL of sterile water before being centrifuged at 2,100 x g for 8 minutes. This centrifugation steps resulted in the formation of a milky white phase between the PBS and salt layers, which contained the live oocysts. Using this methodology, empty and non-viable oocysts fall to the bottom of the tube during centrifugation^53^. The oocysts were carefully pipetted from the interface. Because, the isolated oocysts may contain a carryover from the sodium chloride solution, 9 mL of 1x PBS were added to dilute the mixture and the oocysts were pelleted again at 2,100 x g for 8 minutes. The pipetting/ dilution steps were repeated twice.

### DNA extraction from oocysts and infected cultures

COLO-680N cells infected using the protocol above, were washed twice with 1x PBS, prior to DNA extraction. Samples were taken six hours post infection and subsequently on days 1, 2, 3, 4, 5, 6, 9 and 12.

For COLO-680N cells and epicellular *Cryptosporidium* stages, DNA extraction was performed using the Qiagen DNeasy Blood and Tissue kit (Qiagen, Cat. No 69504) following the manufacturer’s instruction. For *C. parvum* oocysts, samples were collected from cell culture supernatants as described above. Then, DNA extraction was performed using the Omega E.Z.N.A. fungal extraction kit (Cat. No. D3390-1) following the manufacturer’s instructions. Extracted DNA was quantified using a NanoDrop 1000 Spectrophotometer.

### PCR amplification

The PCR reaction contained the following reagents in a 50 μL total volume: 10 μL PCR 5x Flexibuffer, 2 μL MgCl_2_, 1 μL dNTPs (10 mM), 2 μL of forward primer (10 mM), 2μL of reverse primer (10 mM), 31.75 μL H_2_O, 0.25 μL of GoTaq G2 Hot Start Polymerase (Promega, Cat. No. M740A) and 1 μL of the extracted DNA. The list of primers used in the different reactions is shown in the supplementary **Table S2**.

PCR set up conditions were as follows: 1 cycle of 95 °C of initial denaturation for 5 minutes, followed by 36 cycles of 35 seconds denaturation steps at 95 °C, 20 seconds annealing steps at 52 °C and 20 seconds elongation steps at 72 °C. Then a final 10 minutes elongation step at 72°C.

The resulting amplified DNA was visualised on a 1.8 % agarose-TAE gel, stained with ethidium bromide.

### Electron Microscopy (EM)

0.5 cm diameter UV sterilised 200 μm thick aclar film discs (Honeywell international Inc. 5042525) were deposited into a 24-well plate. For each well, 1 mL of COLO-680N cells at a concentration of 2.4 x 10^4^ cells/ml was added. Once the cells had reached 50 to 60 % confluency, the cultures were infected with 50,000 *C. parvum* oocysts, giving an approximate MOI of 0.4. On days 6, 7, 8, 9, 12, the supernatant was removed and cells were washed with 200 mM cacodylate buffer (pH 7.4). The buffer was aspirated and 1 mL of fixative, containing 2.5% Glutaraldehyde in 100 mM cacodylate buffer (pH 7.4), was then added and left at 4°C overnight. Next, each well was washed twice (10 minutes) with cacodylate buffer prior to staining with 1 mL of 1% osmium for 30 minutes at room temperature. The samples were washed and dehydrated through an ethanol series (30 %, 50%, 70 %, 90% and twice 100%) before being embedded in Agar (Agar Scientific) low viscosity resin.

Sections were cut initially with a glass knife and ultrathin sections of 65 nm were cut with a Diatome diamond knife on a RMC-MTXL ultramicrotome. The sections were placed onto 400 mesh uncoated copper grids. The grids were then stained for 45 minutes in 4.5 % aqueous Uranyl Acetate, washed again, and subsequently stained for 7 minutes with Reynold’s lead citrate. Stained grids were then dried for 10 minutes before being visualised with a Jeol 1230 Transmission Electron Microscope at 80kV equipped with a Gatan Multiscan digital camera.

### Fluorescence microscopy

50-60% confluent COLO-680N and HCT-8 cultures were infected with *C. parvum* oocysts in 2-well permanox base chamber slides (Sigma-Aldrich, Cat No C6682), with an approximate MOI of 0.4. At harvesting points, cultures were washed with 1x PBS and then fixed in methanol for 10 minutes at room temperature. Next, the methanol was removed and the cells were permeabilised with 0.002 % Triton-X100 in 1x PBS at room temperature for 30 minutes. Cells were then washed three times prior to incubation for 1 hour with *Cryptosporidium-*specific antibodies (Crypt-a-glo, Waterborne™, dilution 1:10; Sporo-glo, Waterborne™, dilution 1:10; anti-CpClec antibody, dilution 1:60), propidium iodide (500 nM), or *Cryptosporidium*-specific Vicia Villosa lectin^54, 55^ (VVL, 0.5μg/ml).

Cells were washed a further three times with 1x PBS and then mounted with an aqueous mounting medium. This was done using either Fluromount (Sigma-Aldrich, Cat No F4680, with no DAPI) or Fluoroshield (Sigma-Aldrich, Cat No F6057, with DAPI). Slides were visualised by fluorescence microscopy using an Olympus 1X82 or Zeiss Elyra P1 confocal microscope.

### Atomic Force Microscopy (AFM)

1 x 10^5^ oocysts were suspended in 25 μL 1x PBS, pipetted onto a freshly cleaved mica sheet pre-mounted on a magnetic specimen disc (Agar scientific) and left to sediment for an hour at 4°C in a humidified chamber. Oocysts were then fixed for 1h with 25 μL of 5% Glutaraldehyde in 1x PBS. Samples were then washed twice with 1x PBS, once with dionised water and air-dried at room temperature in a dust free area for 2 hours. Samples were then washed twice more with dionised water, left to air-dry again followed by a final drying step using a gentle stream of Nitrogen.

Samples were analysed by Atomic force microscopy (AFM) using a Bruker multimode 8 (Bruker Corporation, Massachusetts) scanning probe microscope with a Nanoscope V controller. The samples were imaged using the ScanAsyst peak force tapping mode, with RTESPA silicon cantilever probes (Bruker Corporation, Massachusetts), which have a nominal spring constant of 40 N/m and a nominal tip radius of 8 nm. Data collection and processing was performed using the Nanoscope software (version 1.40, Bruker Corporation, Massachusetts). Images were scanned over a surface area of at least 10 x 10 μm at a resolution of 2048 x 2048 pixels. The scan rate was 0.2 Hz. Images were processed using the Nanoscope Analysis software (version 1.40, Bruker Corporation, Massachusetts) and custom scripts written in Matlab to remove the sample surface tilt and scanner bow. 3-D representations of the data in height and peak-force error channels were subsequently rendered in Nanoscope Analysis.

Further experimental procedures can be found in the Supplemental material section.

## Acknowledgments

CNM is supported by a GTA studentship from the School of Biosciences, University of Kent and a travel award from the Microbiology Society. LJ is supported by a NIBB BBSRC Proof of concept grant (BioProNET - PoC Nov15 Michaelis) awarded to MM and ADT. ADT is supported by starting funds from the School of Biosciences, University of Kent, and a BBSRC research grant (BB/M009971/1). We would like to thank Prof. Jessica Kissinger for providing us with the synchronisation of infection protocols and Dr. Justin Pachebat, Dr. Kevin Tyler and Prof. Boris Striepen for input/comments on the manuscript.

## Author Contributions

CNM carried out the infections, the cell/parasite counts, the real-time cell analysis, the fluorescence microscopy and analysed all the data. ADT carried out the PCR amplification experiments. IB prepared the samples for all the microscopical techniques, performed imaging, and analysed the data. LJ carried out the infections, fluorescence microscopy and contribute in writing part of the manuscript. BB and WFX acquired and analysed the AFM images. JP prepared and analysed the proteomics and lipidomics data. LY planned and performed cell viability assays and analysed data. MP planned and performed SEM experiments and analysed data. JC planned cell culture experiments, analysed data and wrote methodology. ADT and MM directed research, planned experiments, analysed data, and wrote the manuscript.

## References

1. Kotloff, K. L. et al. Burden and aetiology of diarrhoeal disease in infants and young children in developing countries (the Global Enteric Multicenter Study, GEMS): a prospective, case-control study. Lancet 382, 209–222 (2013).

2. Shirley, D. A., Moonah, S. N. & Kotloff, K. L. Burden of disease from cryptosporidiosis. Curr. Opin. Infect. Dis. 25, 555–563 (2012).

3. Wanyiri, J. W. et al. Cryptosporidiosis in HIV/AIDS patients in Kenya: clinical features, epidemiology, molecular characterization and antibody responses. Am. J. Trop. Med. Hyg. 91, 319–328 (2014).

4. Liu, L. et al. Global, regional, and national causes of child mortality in 2000-13, with projections to inform post-2015 priorities: an updated systematic analysis. Lancet 385, 430–440 (2015).

5. Ryan, U., Paparini, A., Monis, P. & Hijjawi, N. It’s official – *Cryptosporidium* is a gregarine: What are the implications for the water industry? Water Res. 105, 305–313 (2016).

6. Aldeyarbi, H. M. & Karanis, P. The Ultra-Structural Similarities between *Cryptosporidium parvum* and the Gregarines. J. Eukaryot. Microbiol. 63, 79–85 (2016).

7. Caccio, S. M. Molecular epidemiology of human cryptosporidiosis. Parassitologia 47, 185–192 (2005).

8. Wielinga, P. R. et al. Molecular epidemiology of *Cryptosporidium* in humans and cattle in The Netherlands. Int. J. Parasitol. 38, 809–817 (2008).

9. Leoni, F., Amar, C., Nichols, G., Pedraza-Diaz, S. & McLauchlin, J. Genetic analysis of *Cryptosporidium* from 2414 humans with diarrhoea in England between 1985 and 2000. J. Med. Microbiol. 55, 703–707 (2006).

10. Checkley, W. et al. A review of the global burden, novel diagnostics, therapeutics, and vaccine targets for *Cryptosporidium*. Lancet Infect. Dis. 15, 85–94 (2015).

11. Striepen, B. Parasitic infections: Time to tackle cryptosporidiosis. Nature 503, 189–191 (2013).

12. Briggs, A. D., Boxall, N. S., Van Santen, D., Chalmers, R. M. & McCarthy, N. D. Approaches to the detection of very small, common, and easily missed outbreaks that together contribute substantially to human *Cryptosporidium* infection. Epidemiol. Infect. 142, 1869–1876 (2014).

13. Torgerson, P. R. et al. The global burden of foodborne parasitic diseases: an update. Trends Parasitol. 30, 20–26 (2014).

14. Cantey, P. T. et al. Outbreak of cryptosporidiosis associated with a man-made chlorinated lake--Tarrant County, Texas, 2008. J. Environ. Health 75, 14–19 (2012).

15. White, A. C., Jr. Nitazoxanide: an important advance in anti-parasitic therapy. Am. J. Trop. Med. Hyg. 68, 382–383 (2003).

16. Miyamoto, Y. & Eckmann, L. Drug Development Against the Major Diarrhea-Causing Parasites of the Small Intestine, *Cryptosporidium* and *Giardia*. Front. Microbiol. 6, 1208 (2015).

17. Ryan, J. J. & Ray, C. G. in Sherris Medical Microbiology: An introduction to Infectious Diseases 992 (McGraw Hill Professional, 2004).

18. Leitch, G. J. & He, Q. Cryptosporidiosis-an overview. J. Biomed. Res. 25, 1–16 (2012).

19. Sponseller, J. K., Griffiths, J. K. & Tzipori, S. The evolution of respiratory Cryptosporidiosis: evidence for transmission by inhalation. Clin. Microbiol. Rev. 27, 575–586 (2014).

20. Bouzid, M., Hunter, P. R., Chalmers, R. M. & Tyler, K. M. *Cryptosporidium* pathogenicity and virulence. Clin. Microbiol. Rev. 26, 115–134 (2013).

21. Jex, A. R. et al. Cryptic parasite revealed improved prospects for treatment and control of human cryptosporidiosis through advanced technologies. Adv. Parasitol. 77, 141–173 (2011).

22. Karanis, P. & Aldeyarbi, H. M. Evolution of *Cryptosporidium* in vitro culture. Int. J. Parasitol. 41, 1231–1242 (2011).

23. Ryan, U. & Hijjawi, N. New developments in *Cryptosporidium* research. Int. J. Parasitol. 45, 367–373 (2015).

24. Vinayak, S. et al. Genetic modification of the diarrhoeal pathogen *Cryptosporidium parvum*. Nature 523, 477–480 (2015).

25. Morada, M. et al. Continuous culture of *Cryptosporidium parvum* using hollow fiber technology. Int. J. Parasitol. 46, 21–29 (2016).

26. Hijjawi, N. S., Meloni, B. P., Morgan, U. M. & Thompson, R. C. Complete development and long-term maintenance of *Cryptosporidium parvum* human and cattle genotypes in cell culture. Int. J. Parasitol. 31, 1048–1055 (2001).

27. Muller, J. & Hemphill, A. In vitro culture systems for the study of apicomplexan parasites in farm animals. Int. J. Parasitol. 43, 115–124 (2013).

28. Girouard, D., Gallant, J., Akiyoshi, D. E., Nunnari, J. & Tzipori, S. Failure to propagate *Cryptosporidium* spp. in cell-free culture. J. Parasitol. 92, 399–400 (2006).

29. Bouzid, M. et al. A new heterogeneous family of telomerically encoded *Cryptosporidium* proteins. Evol. Appl. 6, 207–217 (2013).

30. Bhalchandra, S., Ludington, J., Coppens, I. & Ward, H. D. Identification and characterization of *Cryptosporidium parvum* Clec, a novel C-type lectin domain-containing mucin-like glycoprotein. Infect. Immun. 81, 3356–3365 (2013).

31. Thompson, R. C. et al. *Cryptosporidium* and cryptosporidiosis. Adv. Parasitol. 59, 77–158 (2005).

32. Fayer, R., Nerad, T., Rall, W., Lindsay, D. S. & Blagburn, B. L. Studies on cryopreservation of *Cryptosporidium parvum*. J. Parasitol. 77, 357–361 (1991).

33. Orita, H. et al. High levels of fatty acid synthase expression in esophageal cancers represent a potential target for therapy. Cancer. Biol. Ther. 10, 549–554 (2010).

34. Zhu, G. et al. Expression and functional characterization of a giant Type I fatty acid synthase (CpFAS1) gene from *Cryptosporidium parvum*. Mol. Biochem. Parasitol. 134, 127–135 (2004).

35. Zhu, G. Current progress in the fatty acid metabolism in *Cryptosporidium parvum*. J. Eukaryot. Microbiol. 51, 381–388 (2004).

36. Povey, J. F. et al. Rapid high-throughput characterisation, classification and selection of recombinant mammalian cell line phenotypes using intact cell MALDI-ToF mass spectrometry fingerprinting and PLS-DA modelling. J. Biotechnol. 184, 84–93 (2014).

37. Dvorak, J. A. et al. The application of the atomic force microscope to studies of medically important protozoan parasites. J. Electron. Microsc. (Tokyo) 49, 429–435 (2000).

38. Tzipori, S. & Widmer, G. A hundred-year retrospective on cryptosporidiosis. Trends Parasitol. 24, 184–189 (2008).

39. Hijjawi, N. Cryptosporidium: new developments in cell culture. Exp. Parasitol. 124, 54–60 (2010).

40. Oberstaller, J., Joseph, S. J. & Kissinger, J. C. Genome-wide upstream motif analysis of *Cryptosporidium parvum* genes clustered by expression profile. BMC Genomics 14, 516-2164-14-516 (2013).

41. Valigurova, A., Hofmannova, L., Koudela, B. & Vavra, J. An ultrastructural comparison of the attachment sites between *Gregarina steini* and *Cryptosporidium muris*. J. Eukaryot. Microbiol. 54, 495–510 (2007).

42. Perkins, M. E., Riojas, Y. A., Wu, T. W. & Le Blancq, S. M. CpABC, a *Cryptosporidium parvum* ATP-binding cassette protein at the host-parasite boundary in intracellular stages. Proc. Natl. Acad. Sci. U. S. A. 96, 5734–5739 (1999).

43. Lemgruber, L. & Lupetti, P. Crystalloid body, refractile body and virus-like particles in Apicomplexa: what is in there? Parasitology 139, 285–293 (2012).

44. Rider, S. D., Jr & Zhu, G. *Cryptosporidium*: genomic and biochemical features. Exp. Parasitol. 124, 2–9 (2010).

45. Slapeta, J. & Keithly, J. S. *Cryptosporidium parvum* mitochondrial-type HSP70 targets homologous and heterologous mitochondria. Eukaryot Cell 3, 483–94 (2004).

46. Keithly, J. S., Langreth, S. G., Buttle, K. F. & Mannella, C. A. Electron tomographic and ultrastructural analysis of the *Cryptosporidium parvum* relict mitochondrion, its associated membranes, and organelles. J. Eukaryot. Microbiol. 52, 132–140 (2005).

47. Ctrnacta, V., Ault, J. G., Stejskal, F. & Keithly, J. S. Localization of pyruvate:NADP+ oxidoreductase in sporozoites of *Cryptosporidium parvum*. J. Eukaryot. Microbiol. 53, 225–231 (2006).

48. Forney, J. R., DeWald, D. B., Yang, S., Speer, C. A. & Healey, M. C. A role for host phosphoinositide 3-kinase and cytoskeletal remodeling during *Cryptosporidium parvum* infection. Infect. Immun. 67, 844–852 (1999).

49. Deng, M., Rutherford, M. S. & Abrahamsen, M. S. Host intestinal epithelial response to *Cryptosporidium parvum*. Adv. Drug Deliv. Rev. 56, 869–884 (2004).

50. Shebl, F. M., Engels, E. A. & Goedert, J. J. Opportunistic intestinal infections and risk of colorectal cancer among people with AIDS. AIDS Res. Hum. Retroviruses 28, 994–999 (2012).

51. Feng, H. et al. Quantitative tracking of *Cryptosporidium* infection in cell culture with CFSE. J. Parasitol. 92, 1350–1354 (2006).

52. Upton, S. J., Tilley, M., Nesterenko, M. V. & Brillhart, D. B. A simple and reliable method of producing in vitro infections of *Cryptosporidium parvum* (Apicomplexa). FEMS Microbiol. Lett. 118, 45–49 (1994).

53. Chen, F. & Huang, K. H. Study on methods for isolation and purification of *Cryptosporidium parvum* oocysts from mouse feces. Zhongguo Ji Sheng Chong Xue Yu Ji Sheng Chong Bing Za Zhi 24, 219–222 (2006).

54. Alcantara Warren, C. et al. Detection of epithelial-cell injury, and quantification of infection, in the HCT-8 organoid model of cryptosporidiosis. J. Infect. Dis. 198, 143–149 (2008).

55. Sharling, L. et al. A screening pipeline for antiparasitic agents targeting cryptosporidium inosine monophosphate dehydrogenase. PLoS Negl Trop. Dis. 4, e794 (2010).

56. Pohlenz, J., Bemrick, W. J., Moon, H. W. & Cheville, N. F. Bovine cryptosporidiosis: a transmission and scanning electron microscopic study of some stages in the life cycle and of the host-parasite relationship. Vet. Pathol. 15, 417–427 (1978).

57. Umemiya, R., Fukuda, M., Fujisaki, K. & Matsui, T. Electron microscopic observation of the invasion process of *Cryptosporidium parvum* in severe combined immunodeficiency mice. J. Parasitol. 91, 1034–1039 (2005).

58. Aldeyarbi, H. M. & Karanis, P. Electron microscopic observation of the early stages of *Cryptosporidium parvum* asexual multiplication and development in in vitro axenic culture. Eur. J. Protistol. 52, 36–44 (2016).

